# Multidimensional single-cell analysis identifies a role of CD2-CD58 interactions for clinical antitumor T cell responses

**DOI:** 10.1101/2022.02.11.479825

**Authors:** Gabrielle Romain, Paolo Strati, Ali Rezvan, Mohsen Fathi, Irfan N Bandey, Jay R T Adolacion, Darren Heeke, Ivan Liadi, Mario L Marques-Piubelli, Luisa M. Solis, Ankit Mahendra, Francisco Vega, Laurence J.N. Cooper, Harjeet Singh, Mike Mattie, Adrian Bot, Sattva Neelapu, Navin Varadarajan

## Abstract

The *in vivo* persistence of adoptively transferred T cells is predictive of anti-tumor response. Identifying functional properties of infused T cells that lead to *in vivo* persistence and tumor eradication has remained elusive. We profiled CD19-specific CAR T cells that comprise the infusion products used to treat large B cell lymphomas using high-throughput single-cell technologies based on Timelapse Imaging Microscopy In Nanowell Grids (TIMING) that integrates killing, cytokine secretion, and transcriptional profiling. Our results show that the directional migration of CD19-specific CAR T cells is correlated with polyfunctionality. We identified that CD2 on T cells is associated with directional migration and that the interaction between CD2 on T cells and CD58 on lymphoma cells accelerates killing and serial killing. Consistent with this, we observed elevated CD58 expression on pre-treatment tumor samples in patients with relapsed or refractory large B cell lymphomas treated with CD19-specific CAR T cell therapy was associated with complete clinical response and survival. These results highlight the importance of studying dynamic T-cell tumor cell interactions in identifying optimal antitumor responses.

**KEY POINTS:**

- Profiling patient infusion products revealed that polyfunctional CAR T cells show directional migration, which is associated with higher CD2 expression
- The ligand for CD2, CD58 is expressed at higher levels in the tumors of lymphoma patients who respond better to CAR T cell treatment

## INTRODUCTION

T cells stably endowed with a genetically encoded chimeric antigen receptor (CAR) targeting CD19 have shown clinical responses in refractory B-lineage leukemias and lymphomas. The potential for durable remissions has motivated the development of CARs targeting antigens other than CD19 for the treatment of hematological and non-hematological malignancies (1–3). While research on engineered T cells has focused on attributes that are common to the entire population of manufactured cells, such as tumor antigen discovery and immunoreceptor design (1, 4, 5), characterization of metrics of individual CAR T cells that define their functional potential, and thus clinical benefit, has not been adequately investigated.

Because of inter- and intra-tumor heterogeneity, technologies that aggregate T-cell biology are unable to accurately capture the complexities of a T-cell product with defined and desired characteristics. For example, it is widely accepted that less differentiated cells have increased proliferative capacity and improved persistence, but individual cells vary in their persistence and functional potential (6–8). Correlative profiling of clinical infusion products from responders and non-responders to CAR T cell therapy has shown that products with less-differentiated, naïve/memory-like T cells are enriched in complete responders whereas signatures associated with T-cell exhaustion (functional impairment) are enriched in non-responders (9–13). While these results have advanced the field of CAR T cell biology, these studies do not directly profile the dynamic interactions between T cells and tumors, and hence cannot identify the precise molecular interactions necessary for optimal antitumor function of CAR T cells.

## RESULTS

We utilized TIMING (14–16) to quantify the dynamic interactions between CAR T cells and tumor cells that lead to polyfunctional T cell responses. We profiled infusion products (IPs) from five axicabtagene ciloleucel products (axi-cel; 19-28z CAR T cells) comprised of predominantly CD8^+^ T cells [54.5-87.2 %] and memory cells [67.1-82.3 %] (**Figure S1**). CAR T cells as effectors, NALM-6 tumor cells as targets, and pre-functionalized beads coated with IFN-γ capture antibody as cytokine sensors, were loaded sequentially onto a nanowell grid array, and the kinetics of killing and end-point cytokine secretion from the same cells was monitored using TIMING (**Figure 1a-b, Supplementary Video 1**). We classified the cells using trained convolution neural networks (CNN), and performed cell segmentation, and tracking using machine learning algorithms (**Figure 1a**). To define the kinetics of interaction between individual T cells and tumor cells that lead to subsequent killing, two interaction parameters, tContact, cumulative duration of conjugation between the first contact to target death; and tDeath, the time between first contact and target apoptosis, were computed (**Figure 1c, Table Supplementary table 1**). We observed that T cells that only secreted IFN-γ (monofunction) exhibited the longest conjugation durations of all functional (killing and/or IFN-γ secretion) T cells (**Figure 1d**). Second, for all killer T cells, tContact was significantly lower than tDeath demonstrating that T cell detachment preceded tumor-cell apoptosis (**Figure 1e**). Collectively these results profiling IPs showed that polyfunctional T cells rapidly terminate synapses with tumor cells whereas non-killer T cells stay attached to tumor cells leading to IFN-γ secretion without termination of the synapse.

**Figure 1.**
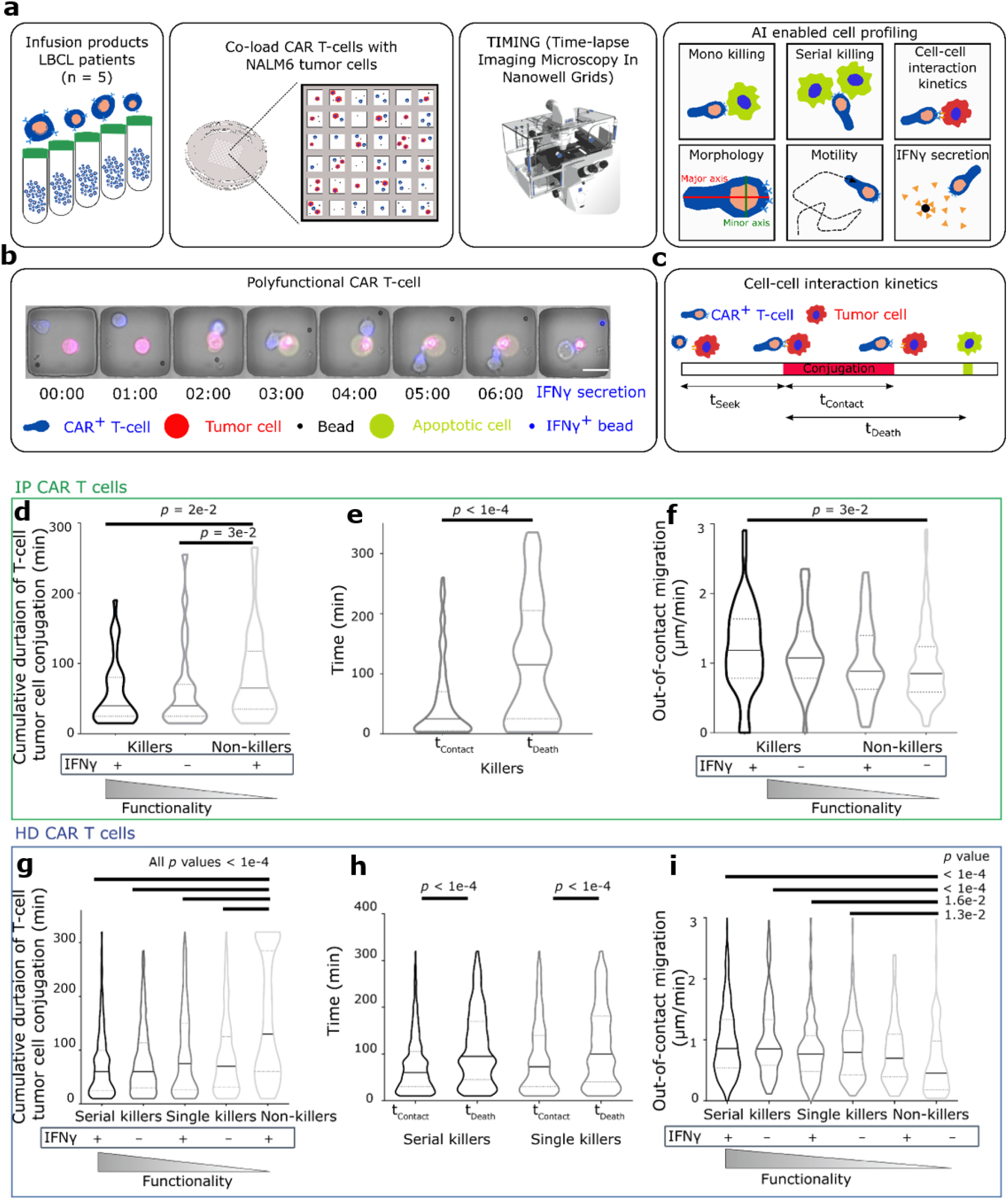
Dynamic single-cell profiling of the polyfunctionality of CAR T-cell infusion products. (a) Schematic overview of the dynamic profiling and image analysis workflow of CAR T cell polyfunctionality using TIMING. We evaluated the interaction between CAR T cells and NALM-6 cells as tumor cells on arrays of nanowells using TIMING. (b) A representative example of a polyfunctional 19-28z T cell that participated in killing and secreted IFN-γ. TIMING is utilized to quantify T-cell intrinsic behavior like directional migration and the kinetics of the interaction leading to induction of apoptosis within tumor cells. After the TIMING assay, the IFN-γ molecules captured onto the beads during TIMING are revealed by using appropriate fluorescently labeled antibodies. Scale bar is 20 μm. (c) Schematic description of kinetic parameters measured in TIMING experiments. (d/g) Cumulative contact duration between effector and tumor cells leading to different functional outcomes. Effector cells that only secrete IFN-γ (monofunctional) exhibited longer contact duration compared to cytolytic cells with or without IFN-γ secretion. The data is aggregated from profiling all five IPs, the bar represents the median and the dotted lines quartiles. (e/h) Comparative assessments of t_Contact_ and t_Death_ for all killer 19-28z T cells. (f/i) Out-of-contact migration of the different functional subsets of 19-28z T cells. All data in (d)-(f) corresponds to E:T of 1:1 and is aggregated from profiling all five IPs (1589 T cells). All data in (g)-(i) corresponds to an E:T of 1:2-5 (to evaluate serial killing) [1178 T cells]. All *p* values for all multiple comparisons were computed using Kruskal-Wallis non-parametric tests and pairwise comparisons using a Mann-Whitney test. The black bar represents the median and the dotted lines denote quartiles.

We next investigated if directional migration (direction of movement is maintained for at least one cell diameter, hereafter migration, **Supplementary Video 2**) can be associated with the efficient exploration of multiple tumor cells T cells leading to polyfunctionality. The search for tumor cells was separated into T cells migrating “out-of-contact” with tumor cells and “in-contact” resulting in conjugation between T cells and tumor cells. Individual killer CAR T cells (had higher out-of-contact migration in comparison to non-killer T cells (**Figure 1f**). These observations were also consistent when measuring in-contact migration during conjugation with the tumor cell, wherein polyfunctional cells had higher migration in comparison to the non-functional cells (**Suppl. Fig. S2**).

Unlike the small numbers of infusion product T cells available for profiling, 19-28z CAR T cells manufactured from healthy donors can be assayed in larger numbers and we next profiled two preclinical 19-28z CAR T cells products (>95% CAR^+^CD8^+^), specifically focusing on serial killing(17). We identified 1,178 nanowells of interest populated with a single live T cell, 2 to 5 NALM-6 tumor cells, and one or more beads. Since every T cell within this subset was incubated with multiple tumor cells, three functional definitions were utilized: single killer cells that killed only one tumor cell, serial killer cells that eliminated at least two tumor cells, and monofunctional cells that did not lyse targets but only secreted IFN-γ. We confirmed that two observations were consistent with the clinical infusion products. First, T cells that only secreted IFN-γ (monofunctional) exhibited the longest conjugation durations of all functional (killing and/or IFN-γ secretion) T cells. These differences were dominated by the presence of a subpopulation of T cells within the IFN-γ monofunctional cells that remained conjugated to the tumor cell for the entire period of observation (**Figure 1g**). Second, for both single-killers and serial killers, t_Contact_ was significantly lower than t_Death_ demonstrating that T cell detachment preceded tumor-cell apoptosis (**Figure 1h**). Serial killer T cells had a lower duration of conjugation to tumor cells in comparison to monokiller T cells, independent of IFNγ secretion (**Suppl. Fig. S3a-b**). Collectively, these results were similar to the IP T cells, and showed that serial killer T cells rapidly terminate synapses with tumor cells whereas non-killer T cells stay attached to tumor cells leading to sustained IFN-γ secretion.

We next confirmed that individual killer CAR T cells (both single and serial killers) had higher out-of-contact migration in comparison to non-killer T cells (**Figure 1i**). Across the different functional subsets of T cells, as defined above, there was an association between decreasing out-of-contact migration and decreasing polyfunctionality (**Suppl. Fig. S4**). These observations were also consistent when measuring in-contact migration during conjugation with the tumor cell, wherein polyfunctional cells and specifically serial killers had higher migration in comparison to the non-functional cells (**Suppl. Fig. S5a-b**). In aggregate, these results demonstrate that migratory T cells are polyfunctional cells capable of killing and serial killing.

To gain an understanding of the mechanism linking migration and function, we integrated dynamic single-cell profiling with transcriptional profiling. We set up a TIMING assay with CAR T cells, without the confounding influence of tumor cells to record basal migration. Single migratory or non-migratory T cells were retrieved (**Figure 2a, Supplementary Video 2 & 3**), and we performed direct quantitative PCR (qPCR)-based amplification since it has higher sensitivity than single-cell RNA-sequencing (17). Seven genes were differentially expressed between the two groups: *CXCR3* and *IL18R1* (chemokine and cytokine receptors); *LAG3, CD244, CD58,* and *CD2* (inhibition/activation receptors) were upregulated in migratory T cells; whereas *CD69* (activation marker) was down-regulated in migratory T cells (**Figure 2b**). Notably, the expression of the *CAR*, *FASLG,* and *GZMB* was no different between the two subsets of T cells (**Suppl. Fig. S6a-c**). *PRF1* showed a trend towards increased expression in the migratory T cells in comparison to the non-migratory T cells, consistent with their increased cytotoxicity but this was not significant (**Suppl. Fig. S6d**). Building upon our recent report (17), these results suggest that the differences in killing capacity between CAR T cells cannot be explained by differences in the abundance of the transcripts of key cytotoxic molecules.

**Figure 2.**
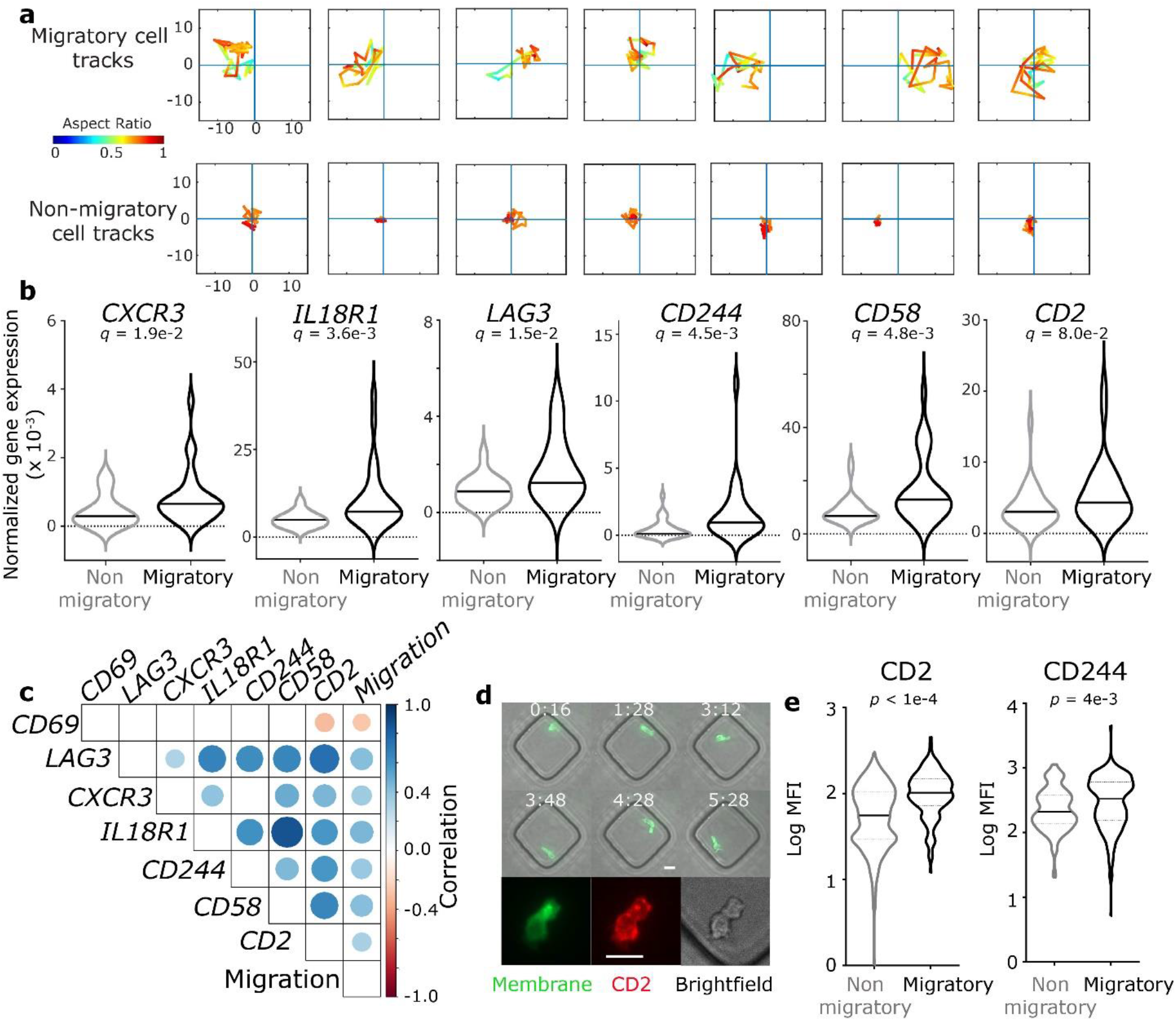
Biomarkers of directional T-cell migration revealed by paired functional and single-cell transcriptional profiling. (a) Representative examples of high and low migration cell tracks during the 3 hours TIMING experiment. X, Y coordinates are shown in microns relative to the initial cell position set to the origin. Color map represents the aspect ratio of cell polarization with red denoting circular cells and increasing shades of green and blue denoting elongated cells. (b) Violin plots illustrating genes differentially expressed between the high and low-migration 19-28z T cells. The genes that are differentially expressed at a false-discovery rate (FDR) q –value < 0.1 are shown. (c) Correlogram illustrating the pairwise correlation coefficients of the transcripts that are significantly linearly correlated with migration. The size of the circle reflects the strength of the correlation and only the significant correlations (p < 0.05) are shown. (d) A representative example of a migratory T cell tracked using TIMING that was subsequently labeled immunofluorescently with the antibody directed against CD2. (e) The differential expression of proteins associated with increased migration of T cells, as determined by immunofluorescent microscopy. Error bars represent 95 % CI and *p* values were computed using Mann-Whitney tests.

To investigate the relationship between the expression of these differentially expressed genes and cellular migration, we computed pairwise correlation coefficients (**Figure 2c**). All seven genes continued to demonstrate a significant correlation to cellular migration, and CD2 transcripts were significantly correlated to all the other genes tested (**Figure 2c**). CD2 and CD244 are members of the CD2 family of proteins and interact with the partner proteins CD58 and CD48 either on other T cells (homotypic) or tumor cells (heterotypic). We quantified the basal migration of CAR T cells in the absence of tumor cells by TIMING and then quantified *a posteriori* the cell surface abundance of CD2 and CD244 by fluorescent immunostaining and microscopy (**Figure 2d**). Comparison of median fluorescence intensity showed that migratory cells had a significantly higher expression of both proteins in comparison to non-migratory cells within the same CAR T-cell populations (**Figure 2e**).

To explore the significance of the interaction of these receptors during interaction with tumor cells, we blocked either CD58 (on NALM-6 targets) or CD2/CD244 (on CAR T cells). We did not block CD48 since leukemic cell lines including NALM-6 do not express this protein(18). We observed that IFNγ secretion was significantly reduced upon blocking either CD2 or CD58 but not CD244 (**Suppl. Fig. S7a**). Next, we performed flow cytometry assays and confirmed that the number of degranulating CAR T cells was also reduced upon blocking the CD58 on tumor cells (**Suppl. Fig. S7b**). To understand the clinical implications of these results, we sought to investigate whether CD58 expression can affect killing mediated by CAR T cells in DLBCL. Accordingly, we expressed CD58 (long isoform) in the HBL-1 DLBCL cell line that lacks CD58 expression (**Figures 3a and 3b**). We tested healthy donor derived 19-28z and 19-41BBz CAR T cells (**Figures 3c and Suppl. Fig. S8**) against parental HBL-1 (CD58^−^) and HBL-1-CD58^+^ cells using TIMING. Flow cytometric analyses confirmed that both sets of T cells had high expression of CD2 (**Figure 3c**). For both 19-28z and 19-41BBz CAR T cells, we observed higher killing against HBL-1-CD58^+^ cells in comparison to the parental HBL-1 cells (**Figure 3d, Supplementary Videos 4 and 5**). Kinetically, this was associated with a shorter duration of synapse formation prior to killing and this ability to form efficient cytolytic synapses is consistent with the known function of CD2 ligation in amplifying the antigen mediated primary signaling (**Figure 3e**)(19). In aggregate, these results show that the interaction between CD2 (T cells) and CD58 (tumor cells) promotes optimal anti-tumor cytolytic functionality.

**Figure 3.**
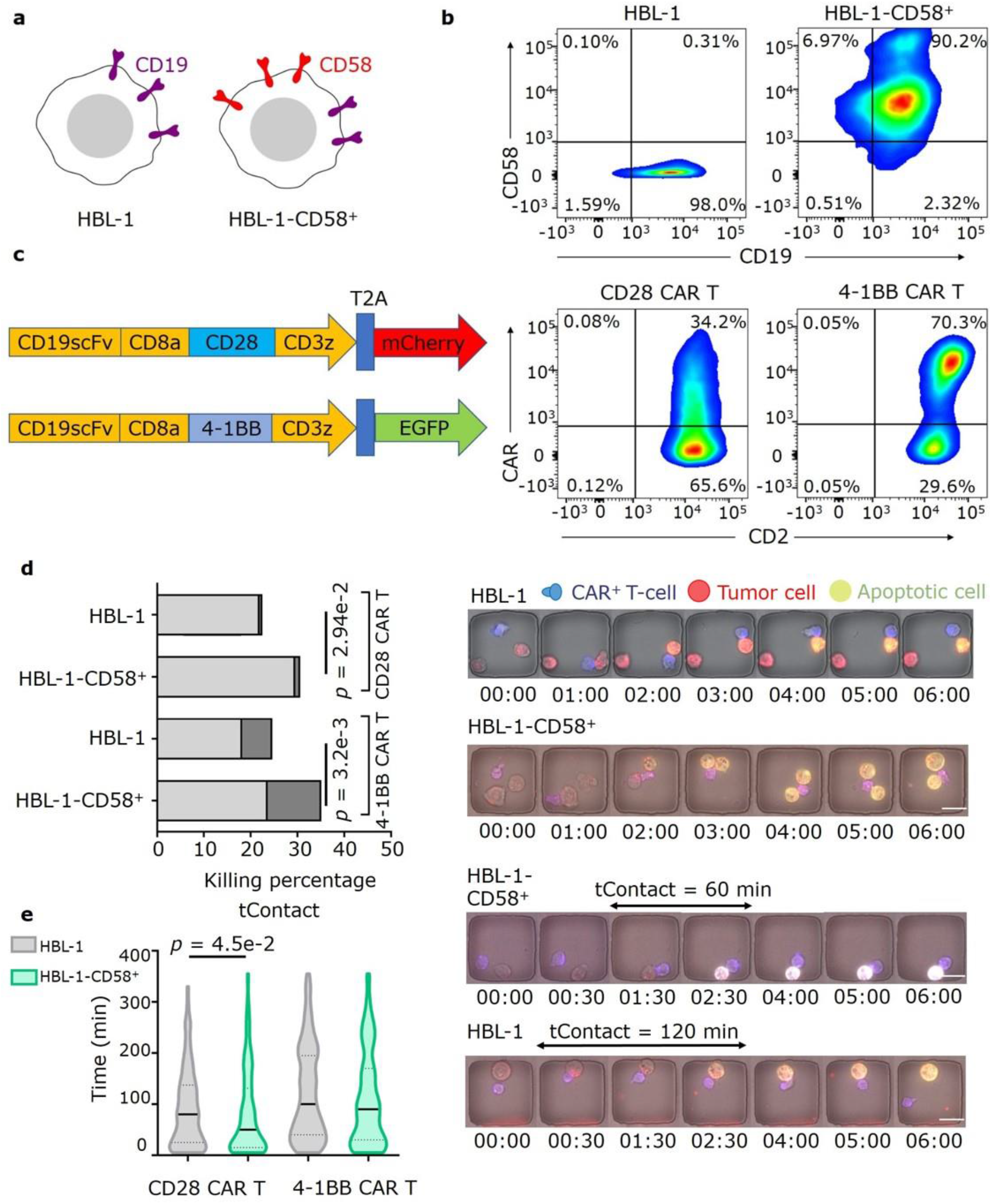
CD58 expression on DLBCL cells enables polyfunctionality of CAR T cells. (a) The DLBCL cell line HBL-1 is deficient in CD58 expression due to homozygous deletion of the CD58 loci. The long form of CD58 was lentivirally transduced into HBL-1 cell lines. (b) Flow cytometric assays demonstrating the relative expression of CD19 and CD58 on HBL-1 and HBL-1-CD58^+^ cell lines. (c) Design of the CAR constructs and CD2 expression in 19-28z and 19-41BBz CAR T cells. (d) T-cell mediated killing of HBL-1 and HBL-1-CD58^+^ cells evaluated using TIMING (E:T 1:1-2). Representative micrographs illustrating monokilling of HBL-1 cells and serial killing of HBL-1-CD58^+^ cells is shown. Scale bar is 20 μm. (e) Kinetic differences in the nature of interaction between CAR T cells and HBL-1 and HBL-1-CD58^+^ cells evaluated using TIMING (E:T 1:1). Representative micrographs illustrating the duration of contact before killing between: 19-28z T cell and HBL-1 cell; 19-28z T cell and HBL-1-CD58^+^ cell. Scale bar is 20 μm. The black bar represents the median and the dotted lines denote quartiles. Pairwise comparisons were performed using Mann-Whitney tests.

We next sought to quantify the impact of CD2 and CD58 expression in patients with B-cell malignancy (large B-cell lymphoma, LBCL) treated with standard-of-care axicabtagene ciloleucel (axi-cel; a 19-28z CAR T cell product). CD2 expression in the infusion product was quantified by scRNA-seq (GSE151511) on CAR T cells (11). Consistent with our preclinical data, the majority of CD3^+^ CAR^+^ T cells (either CD4 or CD8) within both patients achieving complete remission (CR) [59-94 %] and those with progressive disease (PD) [74-98 %] were CD2^+^, and there was no significant difference between the groups (**Suppl. Fig. 9a-b**).

Next, we assessed CD58 expression in tumor cells by chromogenic immunohistochemistry (IHC) in 39 patients with relapsed or refractory LBCL treated with standard-of-care axi-cel, for whom pre-treatment tissue biopsy was available. The baseline clinical characteristics of the patients are described in **Supplementary Table 2**. We observed a complete lack of expression in 12 (31%) cases (**Figure 4a**, left panel; H score = 0) In the remaining 27 (69%) patients, IHC for CD58 showed high intensity with diffuse membranous and cytoplasmatic staining, with a median H-score of 110 [range, 5-300] (**Figure 4a**, right panel).

**Figure 4.**
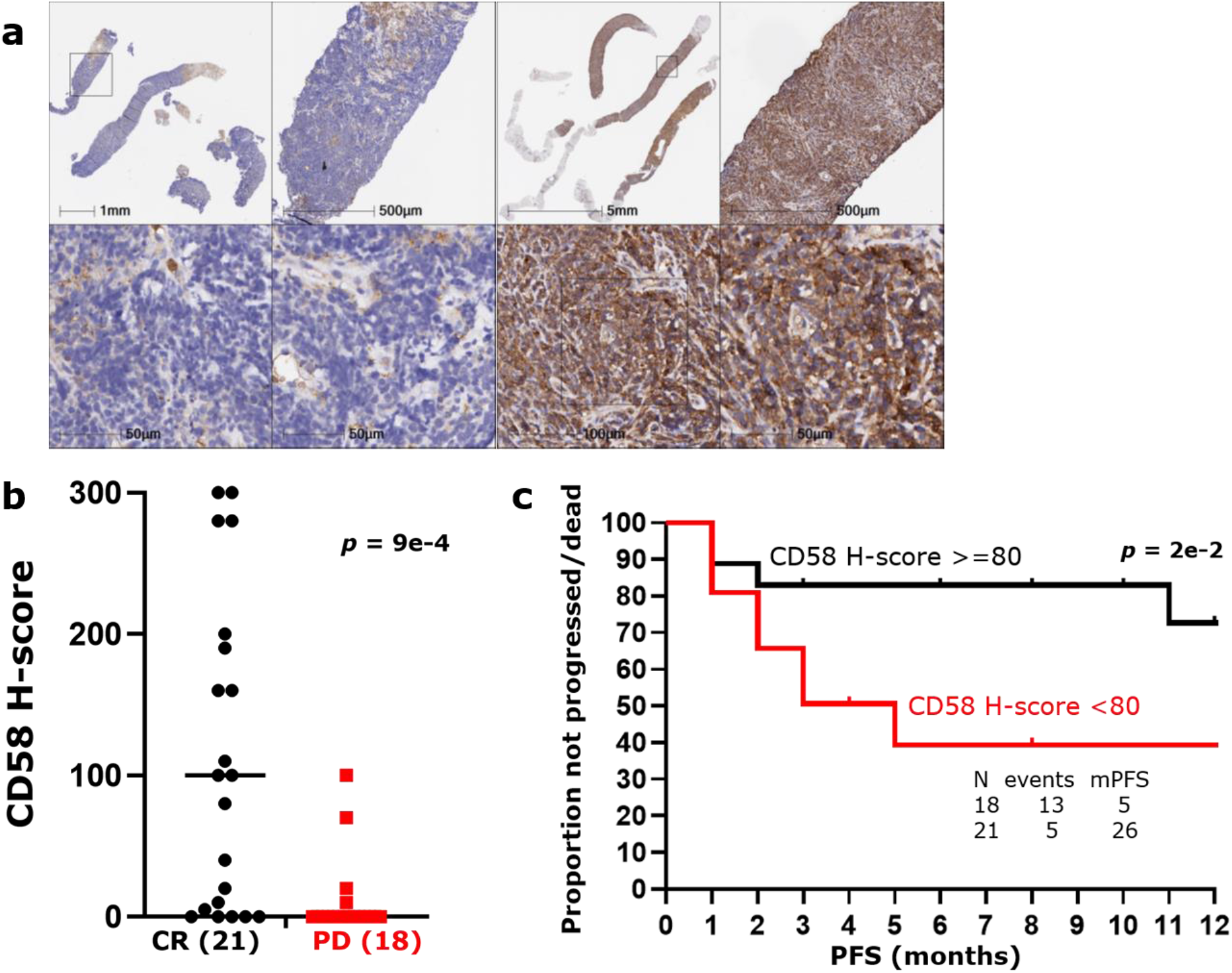
Prognostic role of pre-treatment CD58 expression by IHC in patients with relapsed refractory LBCL treated with axi-cel. (a) CD58 expression by IHC in tissue samples collected before treatment with standard-of-care axi-cel; left panel (negative case): IHC for CD58 shows diffuse negativity of the lymphoma cells with positive internal control (bars show the depth of magnification); right panel (positive cases): IHC for CD58 shows diffuse cytoplasmic and membranous stain with strong intensity in all lymphoma cells (bars show the depth of magnification). (b) CD58 H-score according to day 90 response to standard-of-care axi-cel; bars represent the median; CR, complete remission; PD, progressive disease. *p*-value was computed using a Mann-Whitney test. (c) Progression free survival (PFS) after standard-of-care axi-cel according to pre-treatment CD58 H-score (cut-off=80); N, number; mPFS, median PFS. PFS was calculated using Kaplan-Meier estimates and was compared between subgroups using the log-rank test.

Twenty-one (54%) patients showed CR on positron emission tomography-computed tomography (PET-CT) scan performed on day 90, whereas 18 (46%) were either primary refractory or relapsed/progressed by day 90 after initial response on day 30 (PD). A significantly higher median H-score was observed in patients with ongoing CR at day 90 compared to those with PD (100 vs 7.5) [**Figure 4b**]. After a median follow-up of 12 months (95% confidence interval [CI], 6-18 months), 18 (46%) patients have either progressed and/or died, and median progression-free survival (mPFS) was 14 months (95% CI, 8-20 months). A significantly shorter mPFS was observed when comparing patients with elevated pre-treatment CD58 expression, defined as H-score ≥ 80 (determined by a receiver operating characteristic curve corresponding to a specificity of 0.94), to those with lower expression (26 months vs 5 months, *p* = 2e-2) [**Figure 4c**]. Collectively, these results demonstrate the importance of pre-treatment CD58 expression as a predictive biomarker of response to CAR T cells.

## DISCUSSION

Our data illustrates the impact of directly studying dynamic interactions between T cells and tumor cells in enabling clinically relevant discoveries. Our study of the polyfunctionality of CAR T cells showed that at the single-cell level, non-killer, IFN-γ secreting cells have extended periods of conjugation with tumor cells. The existence of a subpopulation of CAR T cells that remain continuously conjugated to tumor cells due to lack of killing, and leading to IFN-γ secretion, suggests a hypothesis for CAR engineering. Since the functional affinity of the CAR is the primary determinant of the duration of conjugation between the T cell and the tumor cell, engineering CAR designs with altered binding kinetics should identify CARs with preserved polyfunctionality but decreased IFN-γ secretion. This is clinically relevant since chronic IFN-γ secretion is known to be associated with neurotoxicity after CAR T cell infusions (20). Indeed, altered CAR designs with lower affinity for CD19 or humanized CARs with a CD8α hinge have shown robust clinical responses with only minimal serum IFN-γ and associated neurotoxicity (21, 22).

The integrated transcriptional, phenotypic and functional single-cell data demonstrated a correlation between CD2 (LFA-2) and CD58 (LFA-3), and directional migration. Although the positive significance of CD2 is highlighted from pan-cancer studies, it was inferred to merely reflect the presence of infiltrating lymphocytes (23, 24). Our combined functional, transcriptional, and phenotypic data advances the role of CD2-CD58 interactions at the single-cell level and is consistent with independent cell-cell interaction studies probing the genes/proteins essential for cancer immunotherapy using CRISPR-Cas9 screens that mimic loss-of-function mutations involved in resistance to these therapies or in CRISPR-Cas9 screens that identify molecules essential for cytokine secretion (25, 26).

The CD2-CD58 interaction can influence each of the steps that determine the efficacy of infused T cells: migration, function and survival. T cells after infusion need to traffic to the site of the tumor. CD58 is the only T-cell costimulatory molecule expressed by endothelial cells (EC), and the CD2-CD58 interaction between T cells and EC can facilitate recruitment of the circulating T cells to the site of the tumor (27). Once the T cells traffic to the tumor microenvironment, they are likely to encounter tumor cells expressing inhibitory molecules like PD-L1. Within this context, *in vitro* studies support that while PD-1 engagement on T cells can cancel costimulation by CD28, costimulation by CD2 is less sensitive to PD-1 inhibiton (28). Not surprisingly, CD8^+^ tumor-infiltrating lymphocytes in human cancers are characterized by a quantitatively lower expression of CD2, and CD2 mRNA levels in these cells are negatively correlated with exhaustion (19). Independently, it has been shown that CD2 can activate T cells without leading to exhaustion as indicated by the lack of induction of inhibitory receptors including PD-1 (29). T cells are also sensitive to activation-induced cell death (AICD), principally mediated by the ligation of Fas/Fas-L. The binding of CD58 to CD2 on activated T cells protects these T cells from AICD by blocking Fas/Fas-L upregulation, leading to survival and sustained anti-tumor efficacy (30, 31).

From a molecular engineering perspective, CAR structures that incorporate chimeric CD2 signaling endodomains may be compared to CD28 and CD137 with CD3-ζ endodomains as they would be predicted to facilitate anti-tumor responses independent of CD58 on DLBCL (32). From a clinical perspective, patients may be stratified based on CD58 expression and monitored to determine if escape from adoptive cell therapy is accompanied by the expansion of CD58 negative tumor cells (33, 34).

## METHODS

### Human Subjects Statement

All work outlined in this report was performed according to protocols approved by the Institutional Review Boards at the University of Houston and the University of Texas M.D. Anderson Cancer Center.

### Cell lines

Human pre-B cell leukemic line NALM-6 (ATCC) were cultured in T-cell medium (RPMI + 10% FBS) and used as CD19^+^ target cells. HBL-1 DLBCL cell line expressing CD58 was prepared as described previously(35). Parental HBL-1 and HBL-1-CD58^+^ were cultured in IMDM + 10% FBS.

### Manufacture of CAR T cells

We generated CAR T cells as previously described (17). Briefly, we activated PBMCs using OKT3 and anti CD28 antibodies for 48 hours. Next, we transduced the activated T cells with retroviral particles of either CD19R-CD28 or CD19R-4-1BB CAR in RetroNectin coated 24 well plates. The CAR T cells were supplemented with cytokines IL7 and IL15. On day 10, we validated the CAR expression with anti-CAR antibody using flow cytometry. All *in vitro* studies were performed on 10 days old CAR T cells.

### Beads preparation: coating beads with the primary capture antibody

One μL of goat anti-mouse IgG-Fc beads (~2.3 × 10^5^ beads) in solution was washed with 10 μL of PBS and resuspended in 19.6 μL PBS (~0.05% solids). Mouse anti-human IFN-γ (clone 1-D1K) was then added to beads at a final concentration of 10-40 μg/mL and incubated for 30 min at room temperature, followed by washing and resuspension in 100 μL PBS.

### Nanowell array fabrication and cell preparation

We fabricated nanowell arrays for interrogation of effector functions at single-cell level was performed as described previously(36). Approximately 1 million effector cells and target cells were both spun down at 400 × *g* for 5 min followed by labeling with 1 μM PKH67 and PKH26 fluorescent dyes (Sigma-Aldrich) respectively according to the manufacturer’s protocol. Excess unbound dyes were then washed away and cells were re-suspended at ~2 million cells/mL concentration in complete cell-culture media (RPMI + 10% FBS).

### TIMING assays for the multiplex study of effector cytolytic phenotypes and IFN-γ secretion

We loaded capture antibody coated beads and labeled effector and target cells sequentially onto nanowell arrays. Next, detection solution containing Annexin V - Alexa Fluor 647 (AF647) (Life Technologies) (for detection of target apoptosis) was prepared by adding 50 μL of stock solution to 2.5 mL of complete cell-culture media without phenol red. We imaged the nanowell arrays for 6 hours at an interval of 5 minutes using LEICA/ZEISS fluorescent microscope utilizing a 20x 0.80 NA objectives and a scientific CMOS camera (Orca Flash 4.0 v2). At the end of the timelapse acquisition, biotinylated mouse anti-human IFN-γ antibody was added to 2.5 mL cell media at 1:1000 dilution. We subsequently incubated the array for 30 minutes followed by washing and incubation with 5-10 μg/mL Streptavidin conjugated to either R-Phycoerythrin (PE) or Alexa-Fluor-647. We imaged the entire chip to determine the intensity of PE/AF-647 signal on the microbeads and the two datasets were matched using custom informatics algorithms (37).

### Image processing, cell segmentation, cell tracking, and data analytics

Image analysis and cell segmentation/tracking were performed as described previously (16). The pipeline of image processing and cell segmentation ends with statistical data analysis based on the tabular spatiotemporal measurement data generated by the automated segmentation and cell tracking algorithms. We selected nanowells containing single effector and 1-5 tumor cells for further analysis. We then partitioned all these events based on the functionalities of the cells i.e. mono-kill, serial kill, and IFN-γ secretions. A size-exclusion filter based on maximum pixel areas were used to effectively differentiate cells from beads (beads were much smaller than cells). Where specified, cell tracks were represented using MATLAB (Mathworks Inc. MA).

### Gene expression profiling

PKH67 stained CD8^+^ T cells were loaded on a nanowell array, immersed with Annexin V-AF647 containing phenol red-free complete cell-culture medium and imaged for 3 hours using TIMING exactly as described above. After carefully washing the cells on the chip 3 times with cold PBS (4°C), cells were kept at 4^°^C until retrieval. Time-lapse sequences were manually analyzed to identify live cells with high and low migration. The cells were individually collected using an automated micro-manipulating system (CellCelector, ALS) and deposited in nuclease-free microtubes containing 5 μL of 2× CellsDirect buffer and RNase Inhibitor (Invitrogen). Single cell RT-qPCR was then performed using the protocol ADP41 developed by Fluidigm. Ninety-two cells (48 migratory and 44 non-migratory) were assayed, along with bulk samples of 10 and 100 cells, and with no-cell and no-RT controls. The panel of 95 genes included genes relevant to T cell activation, signaling and gene regulation, and was designed and manufactured by Fluidigm D3 AssayDesign(17). Data analysis and processing were performed exactly as described recently(17).

### Flow cytometry-based phenotyping, cytokine secretion, and cytotoxicity assay

For phenotyping, CAR T cells were stained using a panel of human-specific antibodies CD107a (H4A3), CD2 (RPA-2.10), CD58 (1C3), CD244 (2–69), CD62L (DREG-56), CD45RA (HI100), CD45RO (UCHL1), CD95 (DX2), CD3 (SK7), CD27 (L128, MT271), CD28 (L293), CD25 (M-A251), CD127 (HIL-7R-M21), KLRG1 (2F1/KLRG1) CD57 (NK-1). CD4 (OKT4), CD8 (RPA-T8), CD69 (FN50) CCR7 (G043H7), were from BioLegend. The anti-CAR scFv was made in house(38). To confirm CD58 expression on HBL-1 cells, they were stained with human-specific antibodies CD58 (1C3) and CD19 (HIB19) from BD Biosciences. To assay cytotoxicity of the cells at the population level, NALM-6 target cells were stained with PKH26, and co-culture with T cells were set in triplicate at different T cell-to-target cell ratios, with 100,000 T cells per well. Fluorescently labeled anti-CD107a antibodies together with GolgiStop (BD Biosciences) at 0.7 μL/mL was added to the co-culture to stain for degranulating cells. After the assay, cells were washed and stained with Zombie Aqua, antibodies against CD4, CD8, CD3, and CD69, then analyzed by flow cytometry. To measure cytokine release in the supernatants of the co-cultures, IFN-γ and TNF-α were quantified using the MultiCyt® QBeads™ Human PlexScreen (Intellicyt) following the manufacturer’s protocol. For the blocking experiments, CAR T cells were incubated for 24 hours in flat bottom 96 well plates that were pre-coated overnight with 10 μg/mL of purified anti-CD244 (clone C1.7, BioLegend), CD2 (clone LT2, Miltenyi) or anti-CD58 (1C3, BD Biosciences) and rinsed once with complete culture medium before performing the functional assay.

### TIMING and confocal microscopy

CAR T cells were seeded onto nanowell arrays and the migration of the cells monitored using TIMING. At the end of two hours, the cells on the chip were washed carefully three times with cold PBS-5% FBS (4°C). Fluorescently conjugated monoclonal antibodies against CD2, CD244 or CD58 were added to the chip, incubated for 30 min at 4 °C, washed and imaged on a Nikon Ellipse TE confocal microscope fitted with a 60x 0.95NA objective.

### Patient selection and response assessment

This retrospective study was approved by the Institutional Review Board of MD Anderson Cancer Center and conducted in accordance with institutional guidelines and the principles of the Declaration of Helsinki. All patients with relapsed or refractory LBCL treated with standard-of-care axi-cel at MD Anderson Cancer Center, between 01/2018 and 04/2020, were included. The data cut-off for follow-up was 10/20/2020. Response status was determined by Lugano 2014 classification. Pre-treatment tissue biopsies were collected for 21 patients who achieved complete remission after CAR T-cell therapy, and for 21 patients who were primary refractory to CAR T-cell therapy (3 showed crush artifacts, and could not be analyzed)

### CD58 validation, staining, and analysis

Tissue sections of 4 μm thickness were prepared from formalin-fixed paraffin-embedded (FFPE) lymph node and extranodal tissues, and stained using Leica Bond RX automated stainer (Leica Biosystems). The antigen retrieval was performed with Bond ER Solution #1 (Leica Biosystems) equivalent to citrate buffer, pH 6.0 for 20 min at 100℃. Primary antibody against CD58 (CD58/LFA-3 goat polyclonal R&D system,) was used with a concentration of 1:100 (2 μg/ml) and incubated for 15 minutes at room temperature. The antibody was detected using Bond Polymer Refine Detection kit (Leica Biosystems) with diaminobenzidine (DAB) as the chromogen. All the slides were counterstained with hematoxylin, dehydrated, and coverslipped. Tonsils were used as external positive controls and reactive immune cells were used as an internal control. Each case was analyzed using standard microscopy by three pathologists (blinded to response data) and the percentage (10% increments) and intensity (mild, moderate, and intense) of tumor cells staining were evaluated. The data was recorded using H-score system (H-score = [(% mild intensity × 1) + (% moderate intensity × 2) + (% intense intensity × 3)]).

The difference in a continuous variable between patient groups was evaluated by the Mann-Whitney test. Progression-free survival (PFS) was defined as the time from the start of axi-cel infusion to the progression of disease, death, or last follow-up (whichever occurred first). PFS was calculated using Kaplan-Meier estimates and was compared between subgroups using the log-rank test. Receiver operating characteristic (ROC) analysis was used to identify the optimal CD58 H-score for survival analysis (H-score = 80, specificity of 0.94). A *p*-value of <0.05 (two-tailed) was considered statistically significant. Statistical analyses were completed using SPSS 24 (IBM) and Prism 8 (GraphPad).

## ACKNOWLEDGEMENTS

This publication was supported by the NIH (R01CA174385, P30 CA016672); CPRIT (RP180466); MRA Established Investigator Award to NV (509800), Welch Foundation (E1774); NSF (1705464); CDMRP (CA160591); Owens Foundation (NV); The University of Texas MD Anderson Cancer Center B-cell Lymphoma Moonshot (SSN); CRISP Therapeutics and Geron Corporation (FV). We would like to acknowledge the MDACC Flow Cytometry and Cellular Imaging Core facility for the FACS sorting (NCI P30CA16672), UH Seq-n-edit core for RNA-seq, Intel for the loan of computing cluster, and the UH Center for Advanced Computing and Data Systems (CACDS) for high-performance computing facilities. We thank Dr. Riccardo Dalla-Favera for sharing the HBL-1 cell lines.

## AUTHORSHIP CONTRIBUTION

Designed the study: GR, SN, LJNC, HS and NV

Prepared the manuscript: NV, GR, SN, PS, AR, MF and LJNC

Performed experiments: GR, IL, JRTA, IB, HS, PS, AM, MLMP, LMS, AR, MF

Analyzed data: JRTA, GR, NV, IL, PS, MLMP, LMS, AR, MF

Provided patient samples: HS, LJNC, SN, FV, AB, MM, DH

All authors edited and approved the manuscript

