## Supplemental figures and table for "Multidimensional single-cell analysis identifies a role of CD2-CD58 interactions for clinical antitumor T cell responses"

**Table ST1. Glossary of terms used in the paper**

| | |
|---|---|
| CAR | Chimeric Antigen Receptor |
| Directional migration | Motion wherein the direction of movement is maintained for at least one cell diameter |
| 19-28z | CD19-specific CAR construct with a CD8α spacer and CD28 and CD3-ζ endodomains |
| LBCL | Large B-cell lymphoma |
| DLBCL | Diffuse large B-cell lymphoma |
| Axi-cel | Axicabtagene ciloleucel |
| E | Effector CAR T-cell |
| T | Target cell |
| TIMING | Timelapse Imaging Microscopy In Nanowell Grids |
| Monofunction | The ability of a single T-cell to exhibit exactly one function (killing only or IFN-γ secretion only) |
| Polyfunctional | The ability of a single T-cell to kill multiple tumor cells (with or without IFN-γ secretion) or kill exactly one tumor cell and secrete IFN-γ |
| Single-killing | The ability of a single T-cell to kill exactly one target cells |
| Multi-killing | The ability of a single T-cell to kill two or more target cells |
| Single-killer | T cells that kill exactly one tumor cell at an E:T ratio of 1:2-5 |
| Serial killer | T cells that kill at least two tumor cells at an E:T ratio of 1:2-5 |
| Killing efficiency | Description of the kinetics of killing mediated by individual T cells (Please see $t_{Death}$ below) |
| Conjugation | Stable contact between effector cell and target cell lasting > 5 minutes |
| $t_{Death}$ | Time elapsed between first conjugation and tumor cell apoptosis (Annexin V staining positive) |
| $t_{Contact}$ | Cumulative duration of conjugation between synapse formation and $t_{Death}$ |
| AR | The aspect ratio of polarization represented as the ratio of the minor and major axes of the cell fitted to an ellipse |
| $d_{Well}$ | The net displacement of the T-cell centroid, within the nanowell, averaged over 5-minute intervals |

**Table ST2. Patient baseline clinical characteristics collected before initiation of lympho-depleting chemotherapy (day -5)**.

| Patients (N=39) | Median [Range]<br>Number (%) |
|---|---|
| DLBCL/HGBCL, N (%) | 31 (79) |
| Age (years) | 58 [18-84] |
| Male, N (%) | 28 (72) |
| ECOG performance status > 0, N (%) | 31 (79) |
| Ann Arbor Stage III-IV, N (%) | 33 (85) |
| IPI score 3-4, N (%) | 19 (49) |
| C-reactive protein (mg/L) | 29 [0.4-371] |
| Ferritin (mg/L) | 855 [36-9949] |
| Lactate dehydrogenase > ULN, N (%) | 26 (67) |
| Previous therapies (number) | 4 [2-7] |
| Refractory disease, N (%) | 29 (74) |
| Previous autologous SCT, N (%) | 9 (23) |

DLBCL, diffuse large B-cell lymphoma; HGBCL, high-grade B-cell lymphoma; comparator: transformed follicular lymphoma and primary mediastinal B-cell lymphoma; ECOG, Eastern Cooperative Oncology Group; IPI, internal prognostic index; SCT, stem cell transplant; N, number; ULN, upper limit of normal

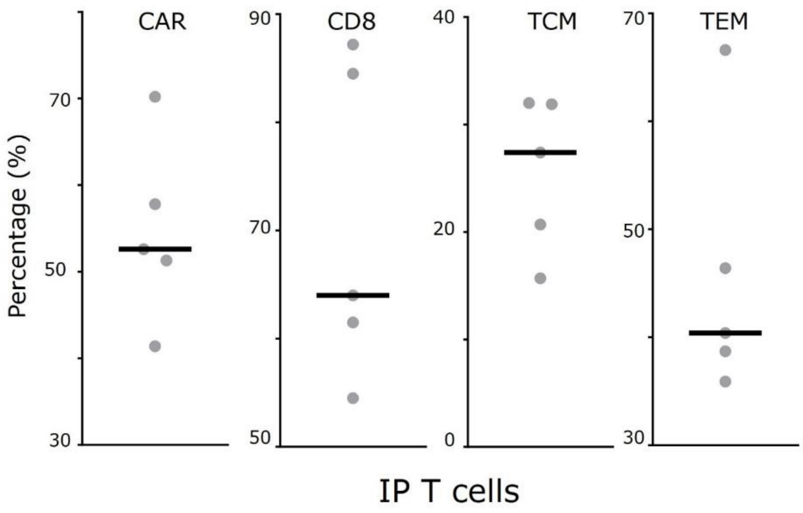


**Supplementary Figure S1**. **Phenotypic characteristics of the five IP CAR T cells profiled using TIMING, determined by flow-cytometry**. TCM and TEM are defined by flow cytometry by immunofluorescent staining of CD3+CD8+CAR+ cells using CD45RA and CD62L.

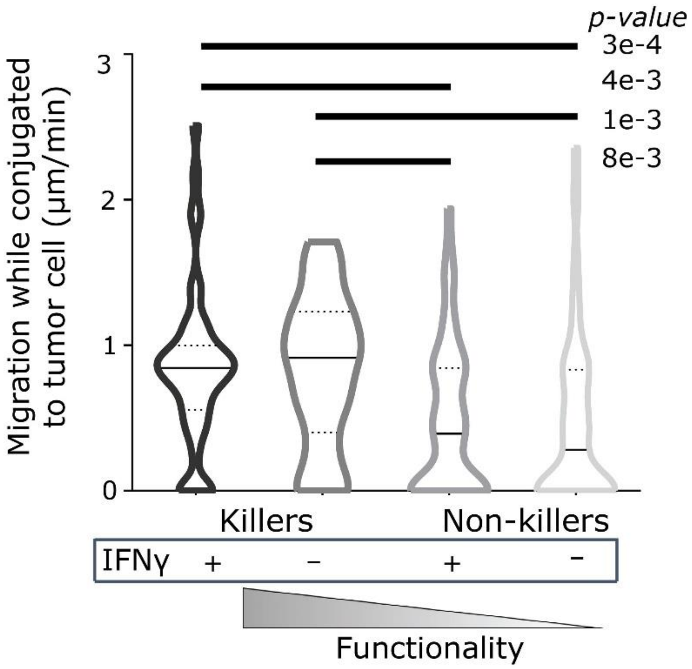


**Supplementary Figure S2. T-cell migration during conjugation to tumor cells is associated with cytolytic activity in IP T cells.**

Effector cells that kill targets, irrespective of whether they secrete IFN-γ, are significantly more migratory compared to non-functional effectors that do not kill or secrete IFN-γ. The bar represents the median and the dotted lines the quartiles. *p* values were determined using Kruskal-Wallis non-parametric test.

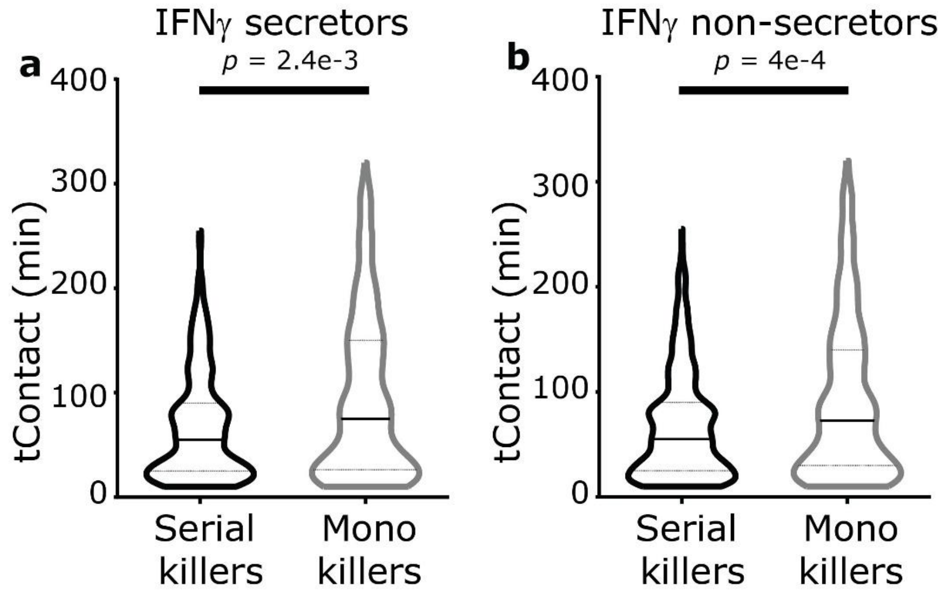


**Supplementary Figure S3. Killer T cells efficiently terminate synapses with tumor cells.**

(a-b) $t_{Contact}$, the cumulative duration of conjugation between T cell and tumor cell before killing, was significantly lower for serial killer 19-28z T cells in comparison to monokiller 19-28z T cells both (a) with, and (b) without IFNγ secretion. For serial killer 19-28z T cells the $t_{Contact}$ first tumor cell killed is shown.

All data were derived from an E:T ratio of 1:2-5. For all panels, *p*-value was determined using a Mann-Whitney test and error bars denote SEM.

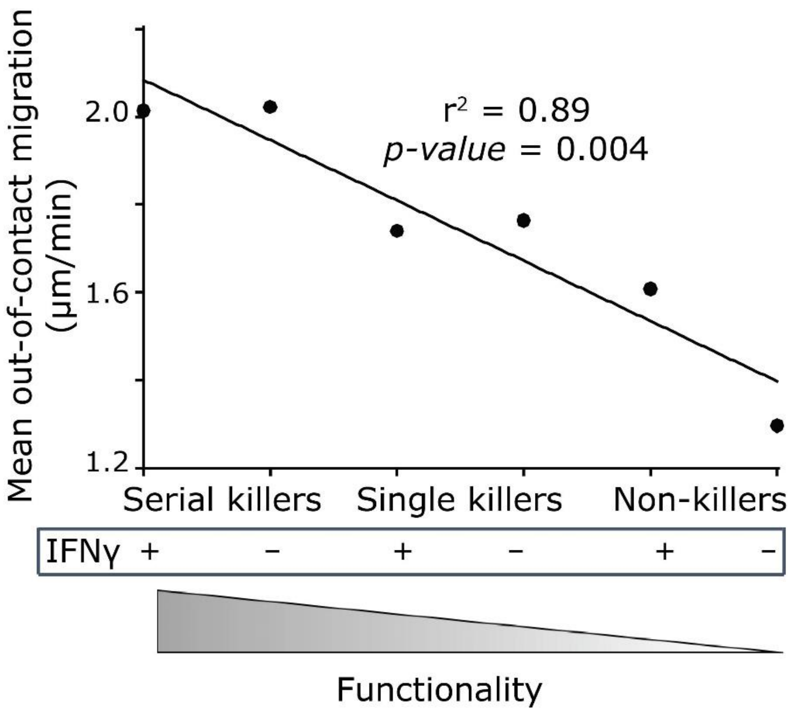


**Supplementary Figure S4. Inverse correlation between polyfunctionality and mean out-of-contact T-cell migration**. All data were derived from an E:T ratio of 1:2-5.

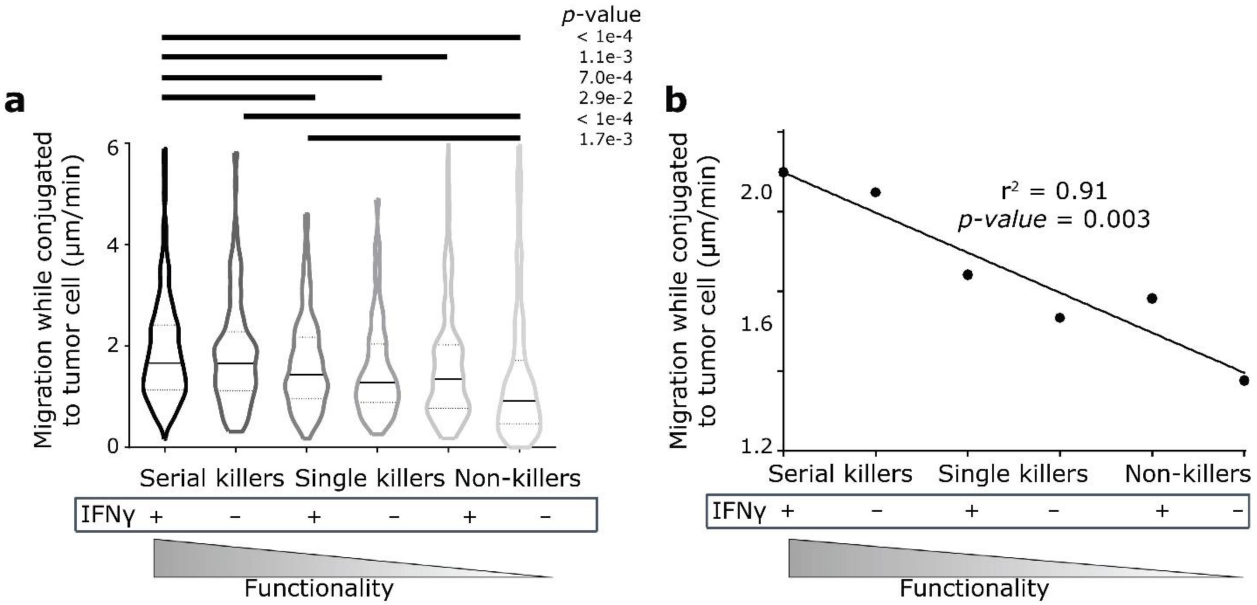


**Supplementary Figure S5. Inverse correlation between polyfunctionality and mean in-contact T-cell migration during conjugation to tumor cells.** At an E:T ratio of 1:2-5, average displacements of effector cells during conjugation of effector with target cells.

(a) Effector cells that kill multiple targets irrespective of whether they secrete IFN-γ, are significantly more migratory compared to non-functional effectors that do not kill or secrete IFN-γ. The bar represents the median and the dotted lines the quartiles. *p* values were determined using Kruskal-Wallis non-parametric test.

(b) Inverse correlation between polyfunctionality and mean T-cell migration during conjugation to tumor cells.

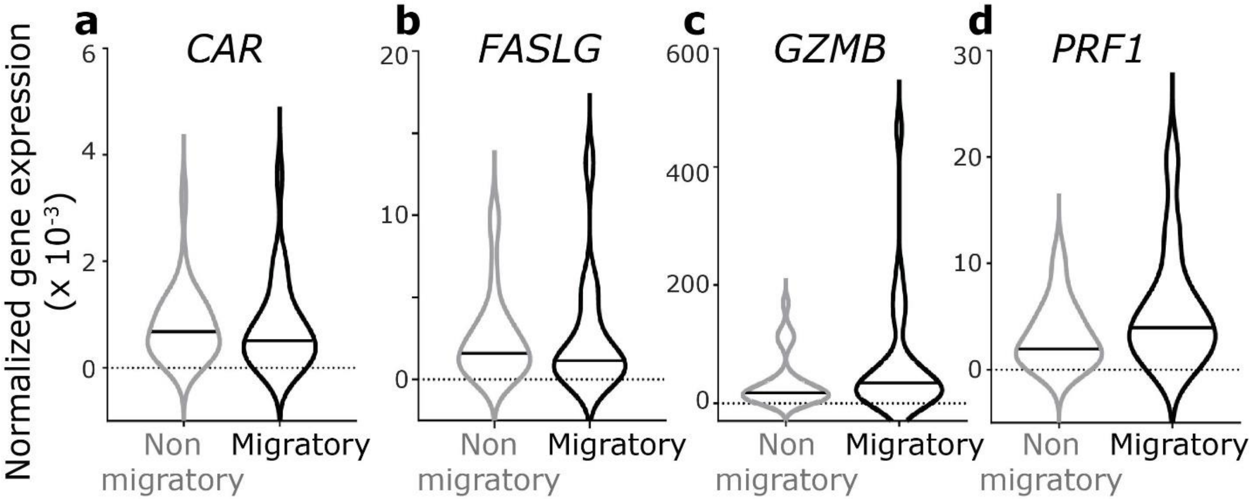


**Supplementary Figure S6. Genes related to cytotoxicity are not differentially expressed between non-migratory and migratory 19-28z T cells.**

(a-d) Violin plots illustrating differences between non-migratory and migratory 19-28z T cells. None of the genes are differentially expressed at a false-discovery rate (FDR) q–value < 0.1.

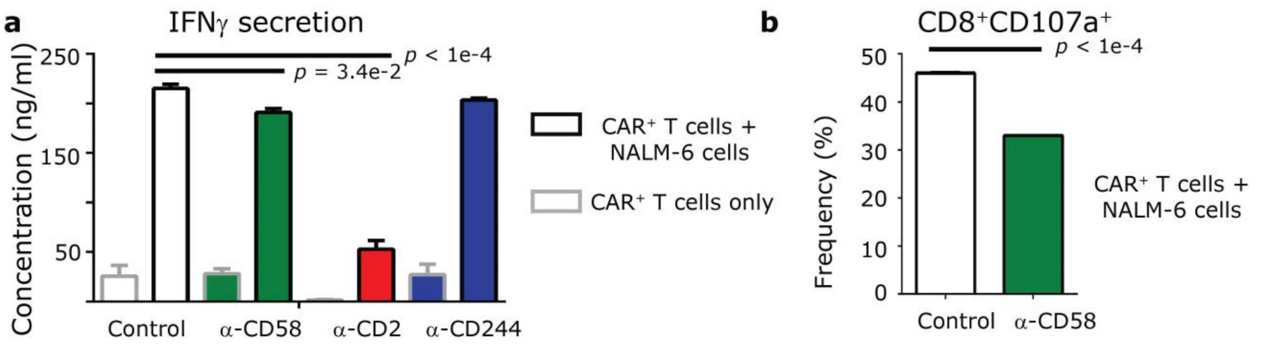


**Supplementary Figure S7. Blocking the interaction between CD2-CD58 negatively impacts CAR T-cell functionality.**

(a) The impact of blocking CD2, CD58, or CD244 determined by measuring cytokine secretion as determined by Intellicyt bead-based assays. These experiments were conducted on a minimum of three donor-derived 19-28z T cells. The error bars represent standard deviation and the statistical test was one-way ANOVA.

(b) Flow cytometric assays enumerating the frequency of degranulating 19-28z T cells upon incubation with tumor cells. These experiments were conducted on a minimum of three donor-derived 19-28z T cells. Error bars represent SEM and the *p*-value was computed using a t-test.

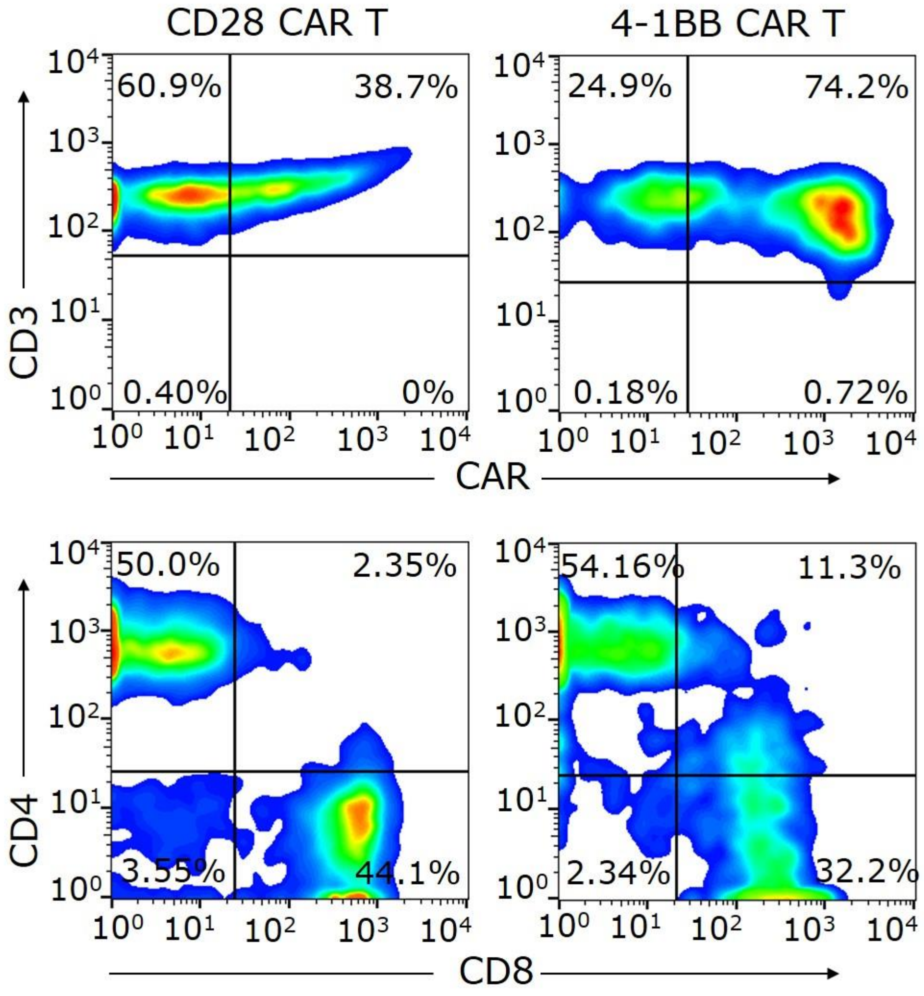


**Supplementary Figure S8. Phenotype of the 19-28z and 19-41BBz CAR T cells derived from healthy donors.**

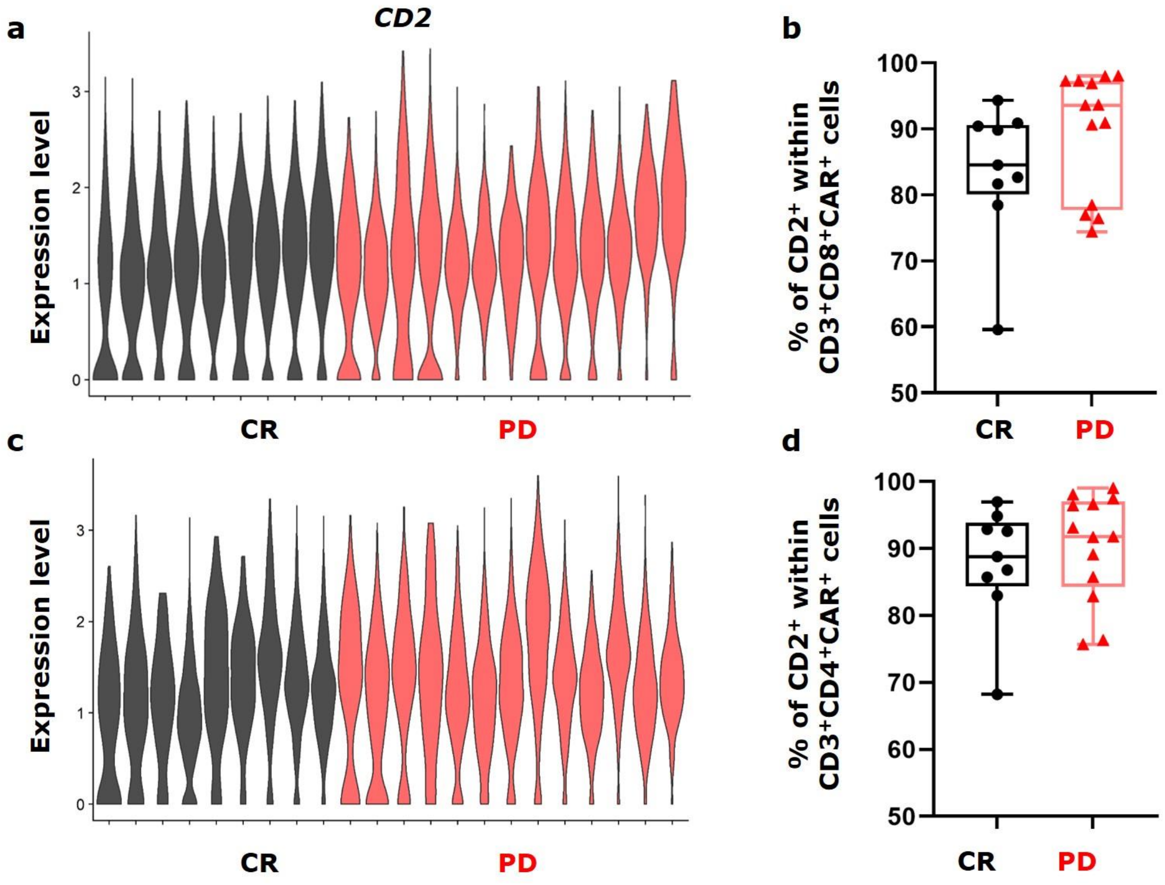


**Supplementary Figure S9.** ***CD2*** **expression in scRNA-seq data derived from 19-28z T cells of clinical infusion products for the treatment of LBCL.**

(a/c) Violin plots of log-transformed gene expression of CD2 within $CD3^+CD8^+CAR^+$ and $CD3^+CD4^+CAR^+$ single cells in twenty-two 19-28z T cell infusion products. The scRNA-seq data is obtained from GSE151511.

(b/d) Comparison of the frequency of $CD2^+$ cells within $CD3^+CD8^+CAR^+$ and $CD3^+CD4^+CAR^+$ cells between complete responders (CR) and patients with progressive disease (PD). Centerline, median. Box limits, upper and lower quartiles. Whiskers, range. Mann-Whitney test *p*-value for both comparisons are not significant.
